# Efficacy of rDiHAVO as a potential vaccine antigen against heartworm infections in a mouse model

**DOI:** 10.64898/2026.09.12.750983

**Authors:** Sucheta Rajaraman, Jhishnu Kannan, Ramaswamy Kalyanasundaram

## Abstract

Dirofilariasis, or heartworm disease, is a life-threatening parasitic infection caused by the nematode parasite *Dirofilaria immitis*. Mosquito vectors in tropical and subtropical regions transmit the infection. Current treatment options are expensive, have a short-term effect, and depend on anthelminthic therapy. There are no licensed vaccines available for preventing heartworm infections. The primary goal of this study was to identify potential vaccine candidates, develop a tetravalent fusion protein (rDiHAVO) vaccine antigen, and evaluate its immunogenicity and efficacy in a mouse model of *D. immitis* infection. Results show that the rDiHAVO is highly immunogenic in mice and confers over 95% protection against a challenge infection. These findings suggest that rDiHAVO is a potential vaccine candidate against Dirofilariasis. Additional testing in dogs can advance this vaccine candidate to clinical trials.

## INTRODUCTION

Dirofilariasis is a vector-borne, zoonotic filarial disease caused by two filarial parasites, *Dirofilaria immitis* and *D. repens*^1–3^. The disease is transmitted by various species of mosquitoes, including *Aedes* sp., *Culex* sp., and *Anopheles* sp., acting as intermediate hosts, facilitating the completion of the parasite life cycle^1, 3^. Infective larvae released into the host during a blood meal develop into mature male and female adults and are present in the pulmonary vessels and heart^3, 4, 5^. Carnivorous animals like dogs, jackals, foxes, and cats are the definitive hosts of the disease, while humans are often the accidental hosts^2^. Dirofilariasis causes a variety of clinical symptoms in dogs, including dyspnea, persistent cough, congestive heart failure, intravascular hemolysis, leading to physical activity intolerance, hemoptysis, pulmonary thromboembolism, ascites, diminished appetite, and weight^4^. The parasite damages the lungs, blood vessels, and heart, resulting in thickening of surrounding tissues, pulmonary artery constriction, and persistent inflammation. Depending on where the worms are, often liver, kidneys, and heart valves are affected^4–7^.

According to the *America*n Heartworm Society 2026 incidence survey, the average number of heartworm cases reported in dogs is increasing across the United States since 2020^7^. Higher incidence is reported in Texas, and new cases are increasing in parts of southwest Colorado, central Washington, northern Idaho, northwest Nevada, and southeast Wyoming. In the USA, about 1.2%-1.4% of dogs are infected with dirofilariasis, with a high incidence among shelter and rescue dogs^8^. Current treatment options include several formulations of melarsomine, which target adult worms or late-stage larvae, and other anthelmintics, mainly Ivermectin, which targets microfilariae^9–11^. Unfortunately, drug-resistant parasites have been reported from all the states in the USA^12, 13^. Doxycycline has been used as a therapy to kill the endosymbiotic bacterium *Wolbachia*, which prevents parasite reproduction^10^. In chronic cases, surgery is the only treatment option for physically removing the worms from the heart^11,14^. If not treated, chronic cases could be fatal. There are no vaccines available to prevent *D. immitis* infection in dogs and cats^1^.

Several previous attempts to develop a vaccine against *D. immitis* suggest that some level of protection can be achieved in laboratory models and dogs^15, 16^. Most importantly, these studies showed that it is possible to develop a protective immune response against *D. immitis* in dogs^15^. Previous reports also identified several antigens as potential candidates for developing a prophylactic vaccine^17–23^. Yet no vaccines are available for preventing dirofilariasis in dogs. Previously, our laboratory and others have screened the genomic library of filarial parasites and identified several potential vaccine candidate antigens^25, 26^. These studies demonstrated that a tetravalent vaccine is more effective than a bivalent or trivalent vaccine against the multicellular filarial nematode parasites^25^. Therefore, in this study, we developed a tetravalent vaccine using four antigens previously shown to have significant vaccine potential. The antigens we used were small heat shock protein^27, 28^ (sHSP), abundant larval transcript 1^29^ (ALT-2), venom allergen^23,30,31^ (VAH), and tropomyosin^32^ (Topo). We combined the nucleotide sequences of the four antigens in tandem to create a tetravalent fusion vaccine (*DiHAVO*). We then expressed rDiHAVO antigen in *E. coli* and evaluated its immunogenicity and vaccine potential against a challenge infection with *D. immitis* infective larvae in a mouse model of dirofilariasis. Our studies show that the rDiHAVO is a promising prophylactic vaccine against *D. immitis* infection.

## MATERIALS AND METHODS

### Antigens used in this study

#### Heat shock protein (DiHSP)

Heat shock proteins are molecular chaperones and are highly conserved across almost all organisms^33^. The conserved α-crystalline domain of small heat shock proteins (HSPs) is essential for their chaperone activity^34^. Small heat shock proteins (HSPs) are increased during the transition from mosquitoes to mammalian hosts, according to a transcriptome study of *B. malayi* L3 larvae^35^. Thus, one essential function of small HSPs is to help the parasite survive in the warmer host. Because of its chaperone function, these elevated amounts of small Hsp12.6 could prevent the parasite’s proteins from being destroyed. Additionally, small HSPs mimic IL-10 and suppress host immune-inflammatory responses that destroy the parasite^36^. Thus, small HSPs aid in the establishment of the parasite in the host and are considered potential vaccine candidates against filarial parasites^35^. HSP also has a potent adjuvant function^37^. The *D. immitis* genome contains sHSP (accession # QHA79233), which was selected to construct the vaccine antigen^27, 38^.

#### Abundant Larval transcript-1 (DiALT)

As the name indicates, the abundant larval transcript (ALT) is the most abundant transcript present in the infective larval stages of all filarial worms, including *D. immitis*^29, 38–40^. Several studies show that ALT can modulate Th2 immune responses in the host and has been successfully demonstrated as a potential vaccine candidate against several filarial parasites^39, 40^. Studies from our laboratory demonstrated that ALT is an excellent vaccine candidate against filarial worms^25^. The *D. immitis* genome contains ALT-1 (accession # MCP9263779), which was selected for use in the construction of the vaccine antigen^19–24^.

#### nDiVA833; Venom allergen (DiVAA)

nDiVA833 shares 13% aa sequence homology with *Ancylostoma ceylanicum* neutrophil inhibitory factor and can block ryanodine receptors^41, 42^. nDiVA833 is critical for parasite development and modulates host immune responses by suppressing T cell activation and T cell proliferation^43–46^. The *D. immitis* genome contains nDiVA833 (VAH) (accession # AAB62535), which was selected for use in constructing the vaccine antigen^19–24^.

#### Tropomyosin (DiTropo)

Tropomyosin is a highly conserved protein essential for maintaining the stability, structure, and rigidity of the parasite’s muscle by regulating actin filament function^32^. Tropomyosin is highly antigenic and has several T cell and B cell epitopes promoting significant cellular and antibody responses^47, 48^. Tropomyosin is secreted by the parasite and is involved in host immune evasion^49^. The *D. immitis* genome contains Tropomyosin (Tropo) (accession # MCP9260322), which was selected to construct the vaccine antigen^19–24^.

#### Tetravalent fusion of all four proteins (rDiHAVO)

The recombinant fusion protein, rDiHAVO, was custom-synthesized at Genscript Biotech (Piscataway, NJ) in *E. coli* host, as 6x His-tagged rDiHAVO protein. The gene and protein sequences of DiHAVO have now been submitted to GenBank (SUB16477902).

### Animals and Parasites

The use of animals in this study was approved by the Institutional Animal Care and Use Committee (IACUC) of the University of Illinois, College of Medicine, Rockford. Twenty-five (25) male eight-week-old BALB/c mice were purchased from Jackson Laboratories (Bar Harbor, ME) and quarantined for 10 days before initiating the experiment. Previous studies showed no significant differences in vaccine-induced immunogenicity and protection between male and female mice^25^. The infective larval stage (L3) of *D. immitis* was obtained from the NIAID/NIH Filariasis Research Reagent Resource Center (University of Georgia, Athens, GA, USA).

### Adjuvant

In this study, we used alum-adsorbed glucopyranosyl lipid (GLA) as the adjuvant (AL019), purchased from the Access to Advanced Health Institute (AAHI, Seattle, WA). GLA is a synthetic TLR4 agonist. Previous studies have shown that AL019 is an excellent adjuvant for filarial recombinant proteins and promotes a balanced Th1/Th2 immune response, which is essential for protection against the multicellular filarial helminth parasites^25^.

### Immunization with rDiHAVO

Mice were divided into three groups of seven (7) mice each. Each mouse in **Group 1** was immunized with 15 μg of rDiHAVO + Al019 (2.5 μg GLA + 50 μg alum). Both *D. immitis* and *B. malayi* filarial parasites share over 70% genomic similarity^50^. Therefore, in this study, we also tested the vaccine potential of a GMP-manufactured r*Bm*HAXT (ΔCys) vaccine developed for human clinical trials against lymphatic filariasis^25, 51^. The vaccine antigens HSP, ALT, and VAH showed significant sequence similarity between *B. malayi* and *D. immitis*^22, 23^. Thus, each mouse in **Group 2** was immunized with 15 μg of GMP-manufactured r*Bm*HAXT (ΔCys) and 10 μg of AL019 (2.5 μg GLA + 50 μg alum). **Group 3** mice served as a placebo control and received 10 μg of the AL019 adjuvant (2.5 μg GLA + 50 μg alum) per mouse. Each mouse received three immunizations of the respective vaccine subcutaneously at two-week intervals.

### Evaluation of Immunogenicity

About 150 μl of blood samples were collected from the submandibular vein of each mouse on day 0, day 14, day 28, and day 42, and serum samples were prepared. Titer of rDiHAVO or r*Bm*HAXT (ΔCys)-specific IgG antibodies was determined in the serum samples using an indirect ELISA as described previously^52^. Briefly, plates were coated with 1 μg of rDiHAVO or r*Bm*HAXT, blocked, and incubated with various dilutions of the sera samples. After washing, horseradish peroxidase (HRP)- conjugated goat anti-mouse IgG F(ab)_2_ (H&L) (Thermo Fisher Scientific, Rockford, IL) was used as the secondary antibody. 1-Step™ Ultra TMB-ELISA substrate (Thermo Fisher Scientific) was used, and the reaction was stopped with 1N H₂SO₄. Absorbance was measured at 450_nm_ using a BioTek Synergy Neo2 microplate reader, with the signal intensity directly correlating with antigen-specific antibody levels.

We also determined the various antigen-specific isotype-specific antibodies using an indirect ELISA as described above. Biotinylated isotype-specific secondary antibodies against mouse IgG1, IgG2a, IgG2b, IgG3, IgA, IgM, and IgE (BioLegend, San Diego, California, U.S), and HRP-conjugated streptavidin (Thermo Fisher Scientific) were used as secondary antibodies. 1-Step™ Ultra TMB- ELISA substrate (Thermo Fisher Scientific) was used to develop the color, and the reaction was stopped with 1N H₂SO₄. The absorbance values directly reflected the levels of antigen-specific antibodies of various isotypes present in the sera.

### Correlates of Protection (CoP) was determined using Antibody-Dependent Cell-mediated Cytotoxicity (ADCC) assay

CoP was determined using an ADCC assay previously described^51, 52^. The assay principle is that IgG antibodies in serum bind to exposed antigens on the parasite. The Fc regions of IgG antibodies bind to Fcγ receptor-bearing cells, such as macrophages, activate them, and initiate parasite killing^53^. Thus, in this assay, we incubated ten (10) infective larvae of *D. immitis* with 50 μl of serum samples and 1 × 10^6 mouse peritoneal cells for 72 hrs. The assay counts the number of dead larvae at the end of incubation and calculates the percent larval death using the following formula

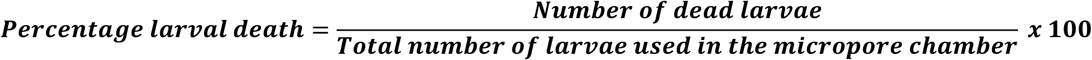

### Challenge studies to determine vaccine-induced protection

All mice were challenged on day 42 using a micropore chamber method as described previously^52^,. Briefly, fifteen (15) infective larvae of *D. immitis* packed into a micropore chamber were surgically implanted into the peritoneum of each mouse. 72 hrs later, the chambers are removed, and the protection was determined as described under ADCC.

### Statistical Analysis

All the data acquired in this study were analyzed using GraphPad Prism 11. Data were expressed as mean ± SD. Significant differences between data were determined using appropriate parametric and non-parametric tests, followed by multiple comparison tests where applicable. Differences with p<0.05 were considered statistically significant with a 95% confidence interval.

## RESULTS

### rDiHAVO vaccine protein has a molecular mass of 75 kDa

About 5 μg of rDiHAVO was separated on a 12% SDS-PAGE gel and stained with Coomassie Brilliant Blue stain (**Figure 1**). 5 µg of BSA was run on an adjacent lane as a molecular mass control. The molecular mass of the rDiHAVO protein was 75 kDa, consistent with the expected value based on the sequences. Several high-molecular-weight bands were visible on the gel, suggesting potential protein aggregation. Based on ImageJ densitometer analysis, nearly 80% of the proteins were at the 75 kDa band.

**Figure 1.**
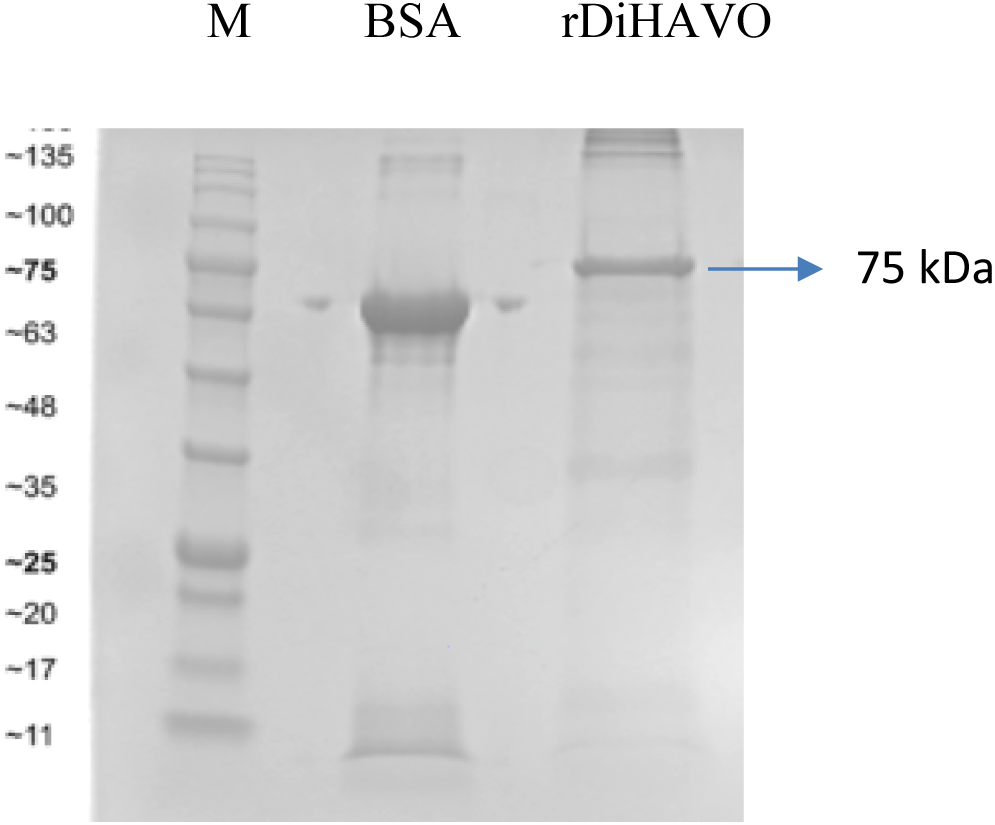
SDS-PAGE analysis of r*Di*HAVO protein supplied by Genscript Biotech. Lane M shows the molecular weight markers; 5 µg of BSA was loaded in an adjacent lane as a molecular mass control. 5 μg of r*Di*HAVO protein was separated on a 12% SDS PAGE gel. A strong 75 kDa band was visible, which is the expected molecular mass of rDiHAVO. A few high-molecular-weight bands were also visible, suggesting possible protein aggregation. Nearly 20% of the protein was aggregated.

#### rDiHAVO immunization induced significant IgG antibody titer

Compared with the control mice that received only adjuvant, all vaccinated mice showed significantly higher (p<0.0001) rDiHAVO-specific IgG titers after three rounds of immunization (**Figure 2**). IgG titers after a single and two immunizations also showed an increase (**Supplementary Figure S1**). The IgG titer (1:5,000) was comparable to the titers obtained after immunization with r*Bm*HAXT (ΔCys). These studies suggested that rDiHAVO is immunogenic. Pre-immune sera samples collected on day 0 did not show any detectable antigen-specific IgG antibodies (**Figure 2**).

**Figure 2.**
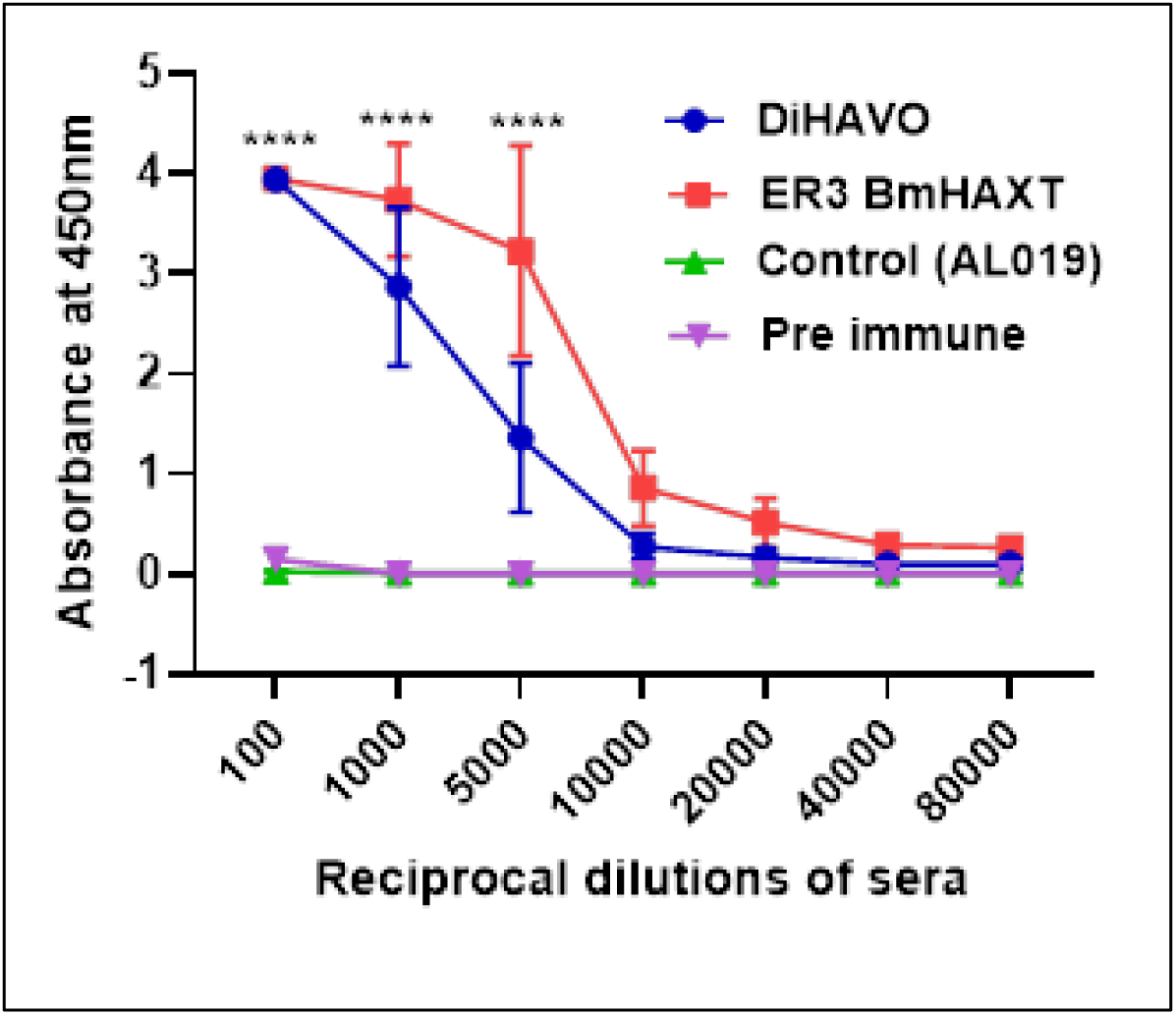
Antigen-specific IgG titers were increased in the sera of immunized mice. IgG titers in the sera of mice immunized three times with the respective antigens were measured by indirect ELISA. Sera samples used in this figure are from mice immunized three times with rDiHAVO or r*Bm*HAXT (ΔCys). Titers up to 1:5,000 were highly significant (****, p<0.0001) compared to the control serum samples. However, there was no significant difference between the rDiHAVO and r*Bm*HAXT (ΔCys) immunized sera samples. Statistical analysis was performed using two-way ANOVA with Tukey’s multiple comparison test. Each data point represents the Mean ± SD (n=7 per group).

The antibody isotype analysis also shows that IgG1, IgG2a, and IgG2b antibody isotypes were significantly (****p<0.0001) elevated in the sera of mice immunized three times with rDiHAVO or r*Bm*HAXT (ΔCys) (**Figure 3**) compared to the control group. These findings suggested that a balanced Th1/Th2 antibody response is generated following the vaccination.

**Figure 3.**
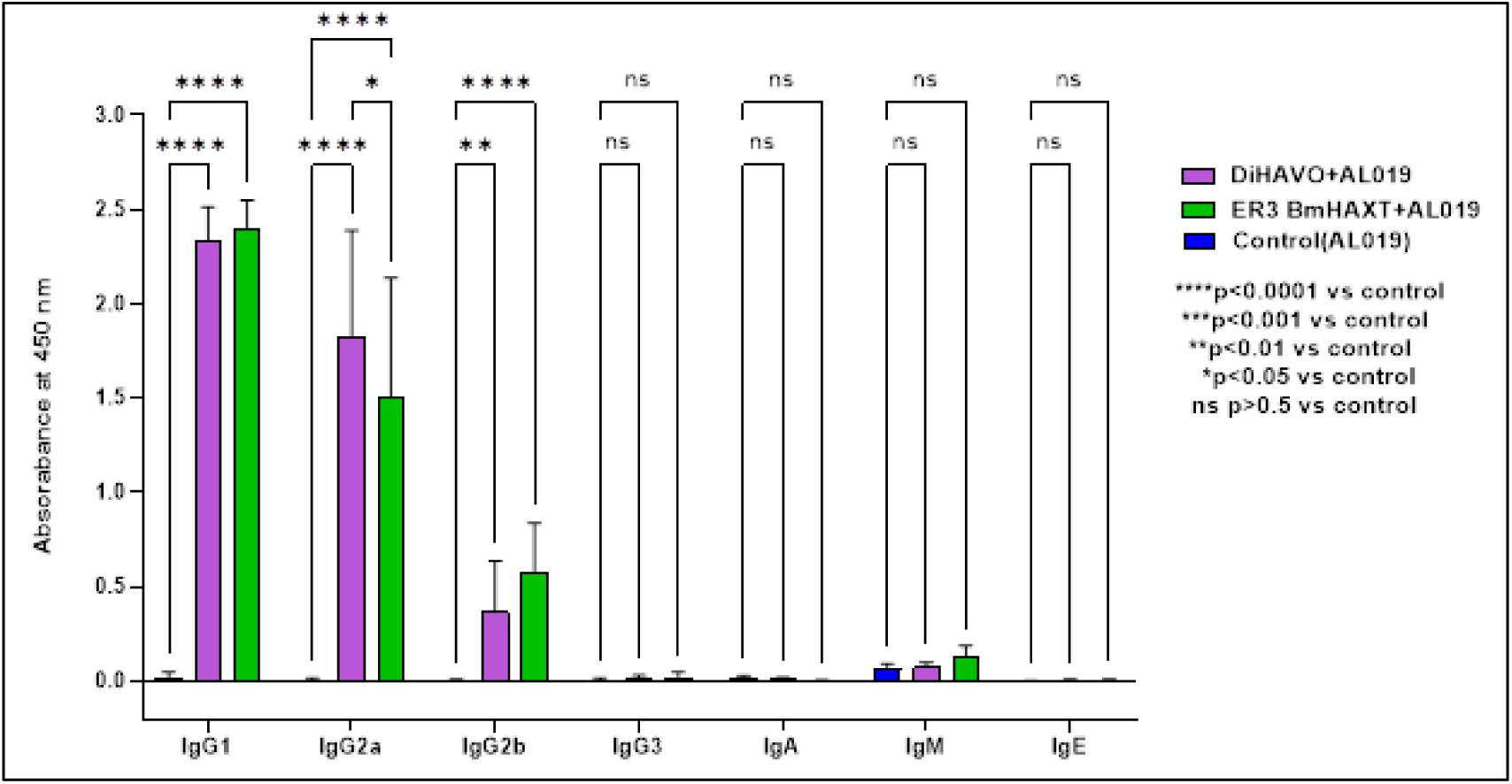
Mice immunized three times with r*Di*HAVO or rBmHAXT (ΔCys) generated high levels of IgG1, IgG2a, and IgG2b antibodies in the sera compared to the AL019 control group. Statistical analysis was performed using two-way ANOVA with Tukey’s multiple comparison test. Each bar represents the Mean ± SD (n = 7 per group).

### Protective antibodies were present in the sera of vaccinated mice

We performed an ADCC assay to demonstrate the correlates of protection (**Figure 4**). Results from this assay showed that the antibodies in the sera of rDiHAVO-immunized mice contributed to 83% killing of infective *D. immitis* larvae. A similar result was observed when sera from r*Bm*HAXT (ΔCys)- immunized mice were used in the ADCC assay, with 87% larval death, compared to controls, which showed 0% larval death (**Figure 4**). These results confirm that immunization with rDiHAVO or r*Bm*HAXT (ΔCys) generates protective antibodies that can mediate ADCC against the infective stages of *D. immitis* in an ADCC assay.

**Figure 4.**
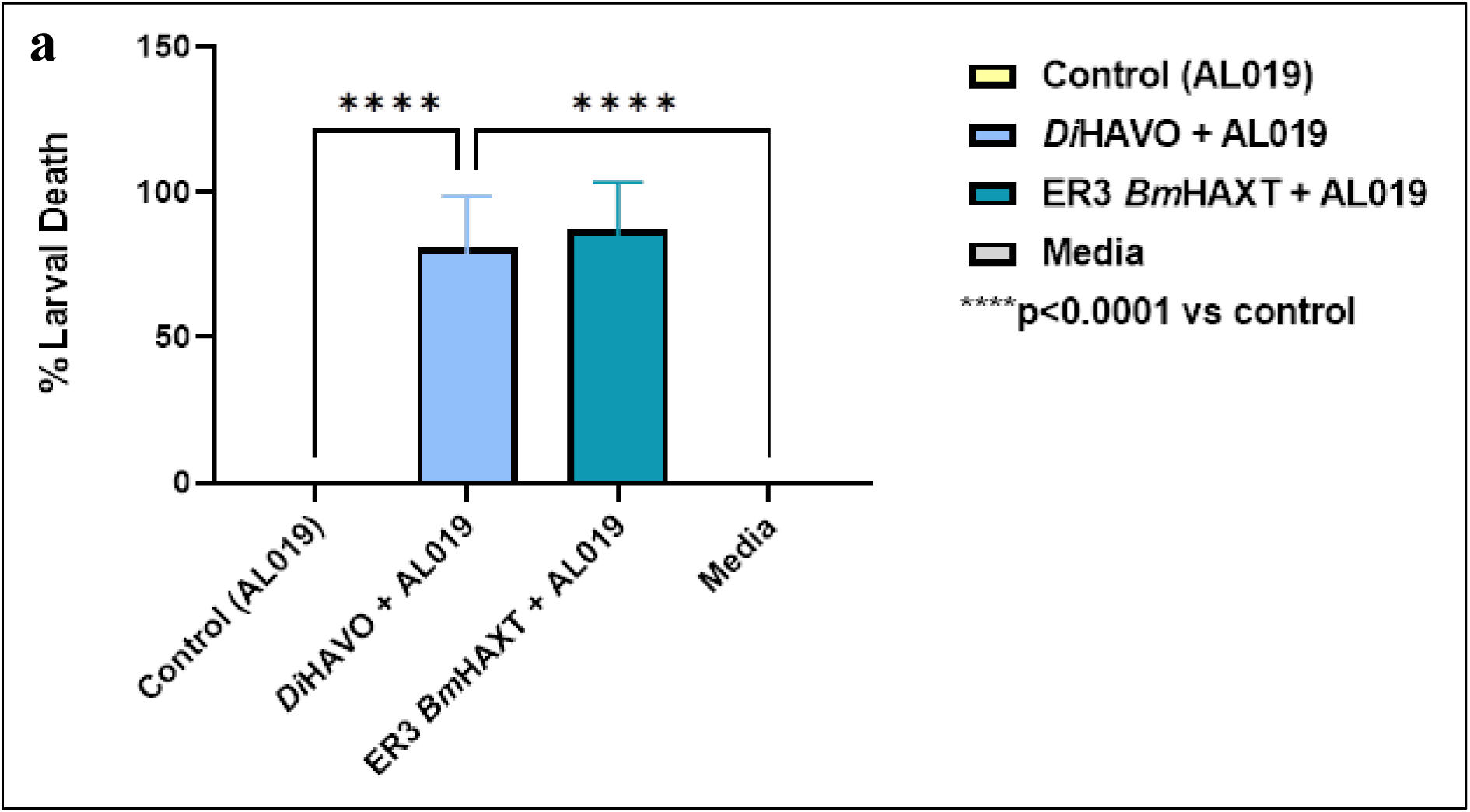
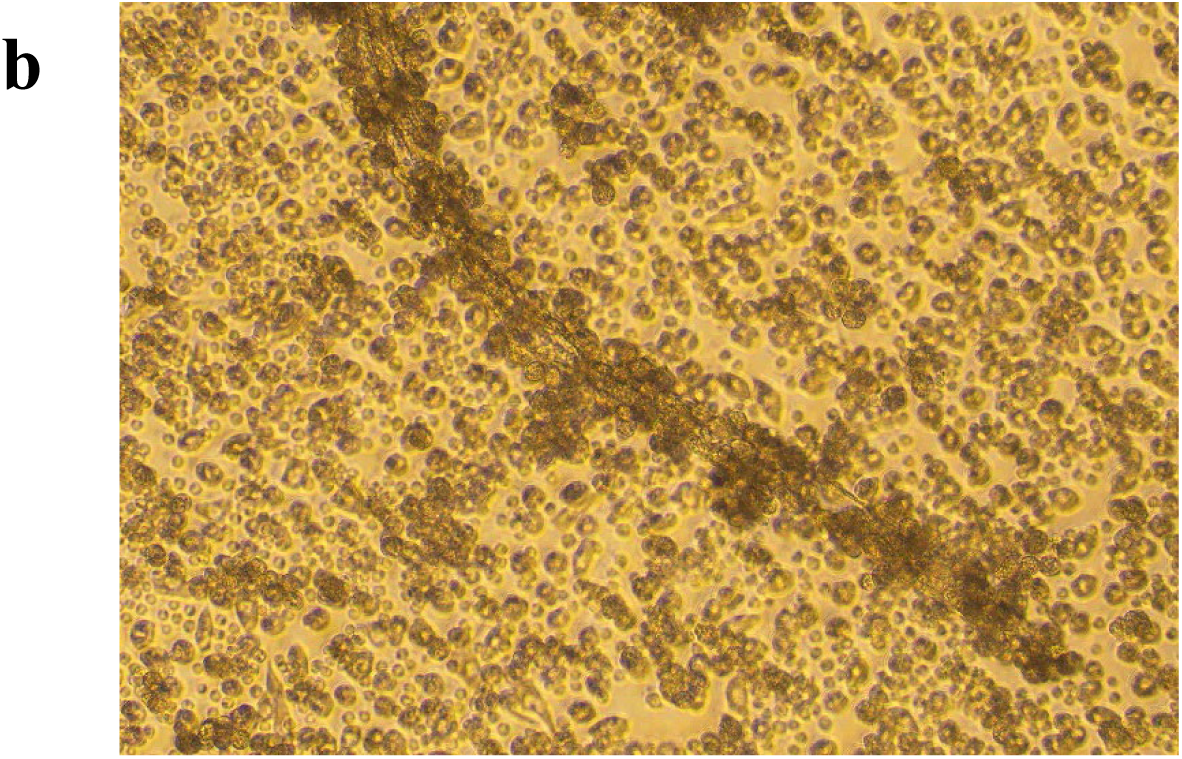
Correlates of Protection: Protective antibodies are present in the sera of immunized mice. (Figure 4a). Sera samples from mice immunized three times with rDiHAVO or r*Bm*HAXT (ΔCys) were incubated with 10-15 infective larvae of *D. immitis* and 1 × 10^6 peritoneal cells for 72 hrs at 37°C. Dead and live larvae were counted in each well. Our data show that 83 ± 1% of the larvae incubated with rDiHAVO sera were dead. The remaining larvae were sluggish and were totally covered with cells, mostly macrophages (Figure 4b). Similar results (87± 2% larval death) were obtained when infective larvae were incubated with sera from mice immunized three times with r*Bm*HAXT (ΔCys) (data not shown). Sera from control mice did not kill the larvae. In these wells, larvae were active and molting, with very few or no cells attached (Figure 4b). These results show that immunization with r*Di*HAVO generates protective antibodies in mice. Statistical significance: ****p<0.0001 vs control (AL019) and media. The statistical difference was determined using one-way ANOVA with Tukey’s multiple comparison test. Each bar represents the Mean ± SD (n = 7 per group).

### Immunization with rDiHAVO confers significant protection against a challenge infection with *D. immitis* infective larvae

Challenge studies demonstrated that vaccination with r*Di*HAVO conferred strong protective immune responses against *D. immitis* infective larvae. About 92% protection was achieved following challenge infection in rDiHAVO-vaccinated animals (**Figure 5**). Out of the seven animals, a couple of vaccinated mice showed 100% protection, suggesting that rDiHAVO is a highly promising vaccine against *D. immitis* infection. Vaccination with r*Bm*HAXT (ΔCys) conferred 86% protection against challenge infection, suggesting that the human r*Bm*HAXT (ΔCys) vaccine is a promising vaccine for *D. immitis* infection in dogs as well.

**Figure 5.**
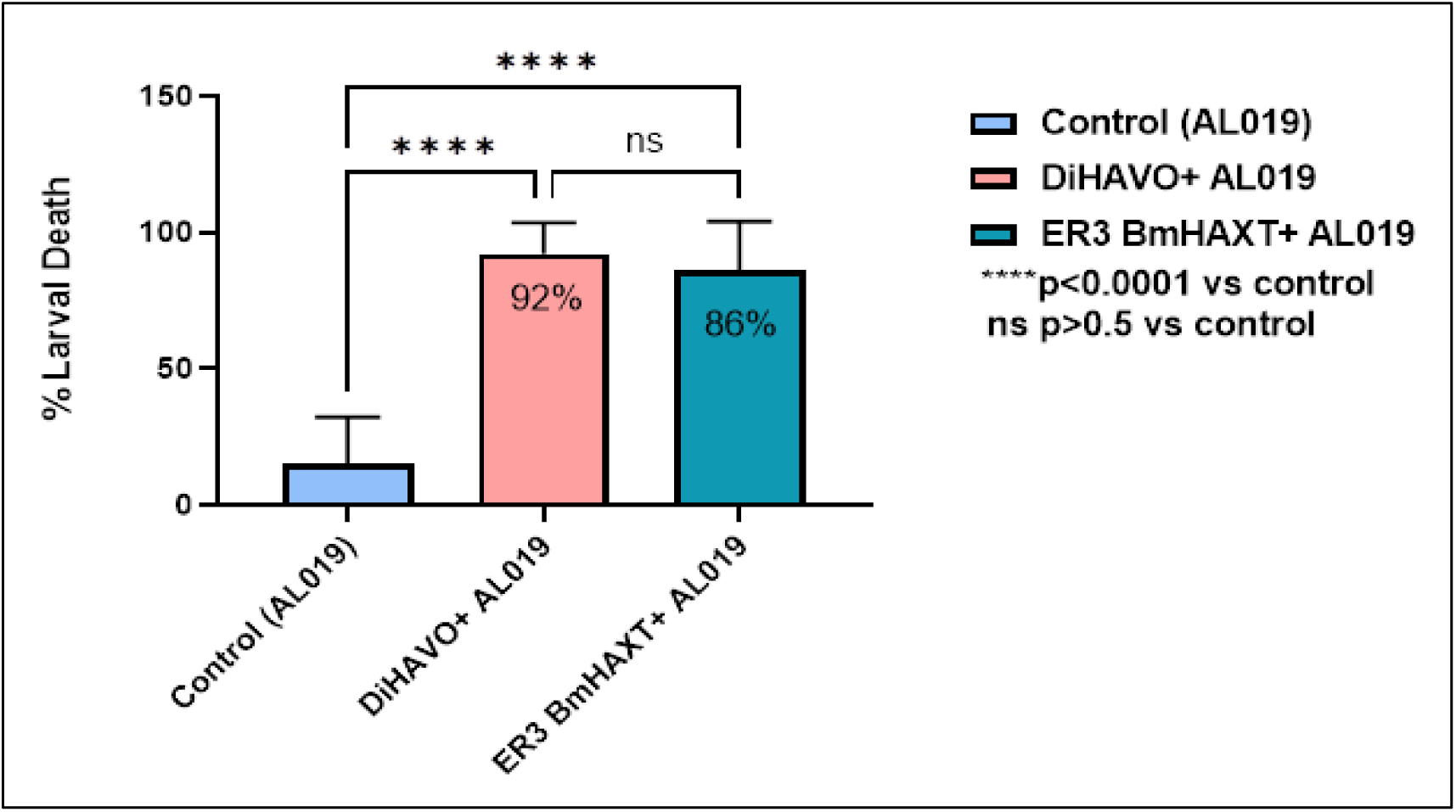
Percent protection after a challenge infection. All mice were challenged with 15 infective larvae of *D. immitis* using a micropore chamber challenge method. The percentage of larval death was determined and used to calculate the percent protection. Results showed that in mice vaccinated with rDiHAVO, the percent protection was 87.7%. compared to controls, which showed 4% larval death. Immunization with r*Bm*HAXT (ΔCys) also conferred significant protection (81.35%), confirming that both rDiHAVO and r*Bm*HAXT (ΔCys) are promising vaccine candidates against *D. immitis* infection. Statistical significance was determined using one-way ANOVA with Tukey’s multiple comparison test. Each bar represents Mean ± SD values (n = 7 per group).

## DISCUSSION

Canine dirofilariasis, caused by the nematode *Dirofilaria immitis*, remains a major parasitic infection in dogs, with significant veterinary and zoonotic implications^1^. This disease, primarily transmitted by mosquitoes, is widespread in temperate and tropical regions, affecting millions of dogs worldwide^2,3^. Current control strategies predominantly rely on macrocyclic lactone (ML) drugs, which prevent larval development and reduce the risk of adult worm establishment^9, 10^. However, issues such as inconsistent administration of preventative treatments, increasing concerns about drug resistance, and limited access to veterinary care in rural and underserved areas have contributed to persistent infection rates and drug resistance^11–13^. Considering these challenges, there is a growing need for novel approaches to control *D. immitis* transmission. In this manuscript, we describe the evaluation of a novel tetravalent vaccine candidate, rDiHAVO, against *D. immitis* in a mouse model. Each candidate antigen in the tetravalent vaccine was selected based on published evidence of their critical role in the parasite’s survival in the host. Immunization with rDiHAVO elicited a strong immune response, characterized by a high titer of antigen-specific IgG antibodies and a balanced Th1/Th2-biased humoral immune response. Using the ADCC assay as a correlate of protection, we demonstrated that these antibodies are protective against infective larvae. Subsequent challenge studies confirmed that the rDiHAVO vaccination confers significant protection against a challenge infection with *D. immitis* infective larvae. We also evaluated the efficacy of an established vaccine candidate for human lymphatic filariasis, r*Bm*HAXT (ΔCys). These studies demonstrated that both rDiHAVO and r*Bm*HAXT (ΔCys) are promising vaccine candidates for *D. immitis* infection.

Efforts to develop a vaccine against canine dirofilariasis have recently gained significant interest due to rising concerns about drug resistance and the limitations of current chemotherapeutic agents^12, 13^. Earlier studies demonstrated that immunization with irradiated *D. immitis* L3 larvae conferred partial protection in dogs, highlighting the immunogenic potential of larval antigens^54^. Building on this, subsequent research used chemically attenuated larvae combined with adjuvants such as Freund’s in Beagle dogs^15^. These studies reported protection rates of up to 77.9%, supporting the feasibility of inducing protective immunity through vaccination. More recently, a project funded by the Morris Animal Foundation at the University of Liverpool has employed genomic and immunological tools to identify conserved antigens across heartworm strains, laying the groundwork for a broadly effective recombinant vaccine^55^. However, currently, there are no vaccines available to protect pets against heartworm infection.

The genome of *D. immitis* is highly similar to that of *Brugia malayi* (the causative agent of Lymphatic filariasis in humans)^50^. Several potential vaccine antigens were identified by genome-wide screening of the *B. malayi* parasite and characterized to develop a tetravalent vaccine for lymphatic filariasis^26, 51^. Leveraging these similarities, we focused on identifying potential vaccine candidates by searching the *D. immitis* genome for antigens that have been successfully used to identify vaccine candidates for *B. malayi* and other major helminth parasites. Major considerations in selecting the final antigens for preparing the vaccine were (i) easy accessibility of the antigens to the host immune system, (ii) antigens that are critical for the survival of the parasite in the host, and (iii) antigens that do not have any homolog in the host^25^. Based on these parameters, we selected four antigens: DiHSP, DiALT, DiVAA, and DiTropo. Previous studies have shown that a combination of multiple antigens is needed for a prophylactic vaccine to combat the multicellular filarial nematode parasites that employ redundant immune evasion strategies to survive in the host^25, 51^.

Since all four antigens were well characterized and had been proven to be excellent vaccine candidates against helminths, we did not evaluate the vaccine potential of each candidate antigen individually. In our previously published studies, we demonstrated that a tetravalent vaccine confers superior protection compared with a monovalent, bivalent, or trivalent vaccine against lymphatic filariasis^25, 51^. We used a mouse model to test our proof of concept. A mouse model has been used to test various parasite vaccines, mainly because the immune responses to these vaccines mimic those in the primary host (in this case, dogs)^19^. Although mice are not permissive hosts for *D. immitis,* infective stage larvae molt to the L4 stage and can be maintained for a few days in immunodeficient mice^56^. Nevertheless, mice are a good immunological model for studying *D. immitis* infection and for obtaining proof-of-concept data to evaluate vaccine immunogenicity and efficacy. Thus, before testing the vaccine in dogs, it is important to evaluate the immunogenicity and correlates of protection of the vaccine candidates in a mouse model.

All mice immunized with rDiHAVO developed high titers of IgG antibodies against rDiHAVO, suggesting that the tetravalent antigen is highly immunogenic. Previous studies using vaccine antigens from helminth parasites demonstrated that the sHSP 12.6^35^, ALT-1^39^, VAH^44^, and Tropomyosin^47–49^ are highly immunogenic proteins. Our ADCC analysis further confirmed that the generated IgG antibodies are protective, and over 83% of infective larvae were killed by the immune sera samples in the presence of cells. We have used a similar assay to determine the correlates of protection in experimental animals (rodents and non-human primates) following vaccination against the lymphatic filariasis worm^51, 52, 57, 58^. Several studies show that a balanced Th1/Th2 immune response is important for achieving prophylactic immunity against filarial parasites ^51, 57, 59^. In this study, based on the isotype of rDiHAVO-specific antibodies, we show that rDiHAVO elicits a balanced Th1/Th2 response, with a slight bias towards the Th2 type (IgG1:IgG2a ratio of 1.41), and that this correlates with protection. A similar immune response was observed when we used the heterologous r*Bm*HAXT antigen, suggesting the potential for using r*Bm*HAXT as a vaccine candidate against *D. immitis*.

Challenge studies with infective larvae of *D. immitis* confirmed that over 87.7% of the larvae failed to establish in mice immunized with rDiHAVO, suggesting that rDiHAVO immunization confers significant protection. Thus, our study demonstrates that rDiHAVO has significant prophylactic potential against heartworm infections. However, there is a need to confirm these observations in a dog model of *D. immitis* infection.

## CONCLUSIONS

In this study, we generated a tetravalent vaccine comprising four antigens selected through bioinformatics analysis and genome-wide comparisons between *B. malayi* and *D. immitis* genes. The recombinant tetravalent fusion protein was custom-synthesized, and its immunogenicity and vaccine potential were evaluated in a mouse model. Our data show that significantly high titers of antigen-specific IgG antibodies were generated after three immunizations with the rDiHAVO vaccine. These IgG antibodies were demonstrated to have significant protective function in an ADCC assay. We also showed that a balanced Th1/Th2 response (IgG1, IgG2a, and IgG2b) was evident following the immunization. Subsequent challenge studies showed that the rDiHAVO vaccination can confer over 87.7% protection in the mouse model. These studies thus demonstrated that rDiHAVO is an excellent prophylactic vaccine against *D. immitis* infection. Additional vaccination trials in dogs will demonstrate the clinical value of this vaccine.

## DATA AVAILABILITY STATEMENT

All data sets presented in this study are included in the article.

## ETHICS STATEMENT

The IACUC committee at the University of Illinois College of Medicine at Rockford reviewed and approved the use of mice and the experimental procedures performed in this study.

## AUTHOR CONTRIBUTIONS

SR and RK planned the experiments. RK designed the vaccine antigens, SR, JK, and RK performed all the mouse studies. SR analyzed the data, and SR and RK drafted the manuscript.

## ACKNOWLEDGMENTS

*Brugia* life cycle stages were obtained through the NIH/NIAID Filarial Research Reagent Resource Center (FR3); morphological voucher specimens are stored at the Harold W. Manter Museum at the University of Nebraska, accession numbers P2021-2032.

## REFERENCES

1. Noack S, Harrington J, Carithers DS, Kaminsky R, Selzer PM. Heartworm disease - Overview, intervention, and industry perspective. Int J Parasitol Drugs Drug Resist. 2021 Aug;16:65–89. doi: 10.1016/j.ijpddr.2021.03.004.

2. McCall JW, Genchi C, Kramer LH, Guerrero J, Venco L. Heartworm disease in animals and humans. Adv Parasitol. 2008;66:193–285. doi: 10.1016/S0065-308X(08)00204-2. PMID: 18486691.

3. CDC, https://www.cdc.gov/dpdx/dirofilariasis/

4. Lemos NMO, Alberigi B, Labarthe N, Knacfuss FB, Baldani CD, da Silva MFA. How does *Dirofilaria immitis* infection impact the health of dogs referred to cardiology care. Braz J Vet Med. 2022 Sep 20;44:e002622. doi: 10.29374/2527-2179.bjvm002622. PMID: 36168657; PMCID: PMC9511925.

5. Kaiser L, Spickard RC, Sparks HV, Williams JF. *Dirofilaria immitis*: alteration of endothelium-dependent relaxation in the in vivo canine femoral artery. Exp. Parasitol. 69, 9–15 (1989).

6. Grauer GF, Culham CA, Cooley AJ, Poff BC, Oberley TD, Brownfield MS, Grieve RB. Clinicopathologic and histologic evaluation of *Dirofilaria immitis*-induced nephropathy in dogs. Am J Trop Med Hyg. 1987 Nov;37(3):588–96. doi: 10.4269/ajtmh.1987.37.588.

7 American Heartworm Society, 2025 Heartworm Incidence. https://www.heartwormsociety.org/veterinary-resources/

8. Drake J, Parrish RS. Dog importation and changes in heartworm prevalence in Colorado 2013-2017. Parasit Vectors. 2019 May 6;12(1):207. doi: 10.1186/s13071-019-3473-0.

9. Dirofilariasis -an overview | ScienceDirect Topics. https://www.sciencedirect.com/topics/medicine-and-dentistry/dirofilariasis.

10. Moorhead AR, Evans CC, Sakamoto K, Dzimianski MT, Mansour A, DiCosty U, Fricks C, McCall S, Carson B, Nelson CT, McCall JW. Effects of doxycycline dose rate and pre-adulticide wait period on heartworm-associated pathology and adult worm mass. Parasit Vectors. 2023 Jul 25;16(1):251. doi: 10.1186/s13071-023-05858-2. PMID: 37491306; PMCID: PMC10369763.

11. Ames MK. Heartworm disease in dogs, cats, and ferrets. 2025. https://www.msdvetmanual.com/circulatory-system/heartworm-disease/heartworm-disease-in-dogs-cats-and-ferrets

12. Prichard RK. Macrocyclic lactone resistance in *Dirofilaria immitis*: risks for prevention of heartworm disease. Int J Parasitol. 2021 Dec;51(13-14):1121–1132. doi: 10.1016/j.ijpara.2021.08.006.

13. Neff E, Evans CC, Jimenez Castro PD, Kaplan RM, Dharmarajan G. Drug Resistance in Filarial Parasites Does Not Affect Mosquito Vectorial Capacity. Pathogens. 2020 Dec 22;10(1):2. doi: 10.3390/pathogens10010002.

14. Lee SG, Moon HS, Hyun C. Percutaneous heartworm removal from dogs with severe heartworm (*Dirofilaria immitis*) infestation. J Vet Sci. 2008 Jun;9(2):197–202. doi: 10.4142/jvs.2008.9.2.197. PMID: 18487942; PMCID: PMC2839098.

15. Grieve RB, Abraham D, Mika-Grieve M, Seibert BP. Induction of protective immunity in dogs to infection with *Dirofilaria immitis* using chemically-abbreviated infections. Am J Trop Med Hyg. 1988 Oct;39(4):373–9. doi: 10.4269/ajtmh.1988.39.373.

16. Anthony RM, Rutitzky LI, Urban JF Jr, Stadecker MJ, Gause WC. Protective immune mechanisms in helminth infection. Nat Rev Immunol. 2007 Dec;7(12):975–87. doi: 10.1038/nri2199. PMID: 18007680; PMCID: PMC2258092.

17. Grieve RB, Frank GR, Mika-Grieve M, Culpepper JA, Mok M. Identification of *Dirofilaria immitis* larval antigens with immunoprophylactic potential using sera from immune dogs. J Immunol. 1992 Apr 15;148(8):2511–5. PMID: 1560206.

18. Marcos-Atxutegi C, Kramer LH, Fernandez I, Simoncini L, Genchi M, Prieto G, Simón F. Th1 response in BALB/c mice immunized with *Dirofilaria immitis* soluble antigens: a possible role for Wolbachia?, Vet Parasitol. 2003, 112(1–2): 117–130, 10.1016/S0304-4017(02)00419-3.

19. Ibrahim MS, Tamashiro WK, Moraga DA, Scott AL. Antigen shedding from the surface of the infective stage larvae of *Dirofilaria immitis*. Parasitology. 1989 Aug;99 Pt 1:89–97. doi: 10.1017/s0031182000061060.

20. Scott AL, Diala C, Moraga DA, Ibrahim MS, Redding L, Tamashiro WK. *Dirofilaria immitis*: biochemical and immunological characterization of the surface antigens from adult parasites. Exp Parasitol. 1988 Dec;67(2):307–23. doi: 10.1016/0014-4894(88)90078-1.

21. Geary TG. New paradigms in research on *Dirofilaria immitis*. Parasit Vectors. 2023 Jul 21;16(1):247. doi: 10.1186/s13071-023-05762-9.

22. Frank GR, Grieve RB. Purification and characterization of three larval excretory-secretory proteins of *Dirofilaria immitis*. Mol Biochem Parasitol. 1996 Jan;75(2):221–9. doi: 10.1016/0166-6851(95)02533-2. PMID: 8992320.

23. Godel C, Kumar S, Koutsovoulos G, Ludin P, Nilsson D, Comandatore F, Wrobel N, Thompson M, Schmid CD, Goto S, Bringaud F, Wolstenholme A, Bandi C, Epe C, Kaminsky R, Blaxter M, Mäser P. The genome of the heartworm, *Dirofilaria immitis*, reveals drug and vaccine targets. FASEB J. 2012 Nov;26(11):4650–61. doi: 10.1096/fj.12-205096.

24. Bennuru S, O’Connell EM, Drame PM, Nutman TB. Mining Filarial Genomes for Diagnostic and Therapeutic Targets. Trends Parasitol. 2018 Jan;34(1):80–90. doi: 10.1016/j.pt.2017.09.003.

25. Kalyanasundaram R, Khatri V, Chauhan N. Advances in Vaccine Development for Human Lymphatic Filariasis. Trends Parasitol. 2020 Feb;36(2):195–205. doi: 10.1016/j.pt.2019.11.005.

26. Gnanasekar M, Rao KV, He YX, Mishra PK, Nutman TB, Kaliraj P, Ramaswamy K. Novel phage display-based subtractive screening to identify vaccine candidates of *Brugia malayi*. Infect Immun. 2004 Aug;72(8):4707–15. doi: 10.1128/IAI.72.8.4707-4715.2004. PMID: 15271932; PMCID: PMC470678.

27. Lillibridge CD, Rudin W, Philipp MT. *Dirofilaria immitis*: ultrastructural localization, molecular characterization, and analysis of the expression of p27, a small heat shock protein homolog of nematodes. Exp. Parasitol. 83, 30–45 (1996).

28. Haslbeck M, Franzmann T, Weinfurtner D, Buchner J. Some like it hot: the structure and function of small heat-shock proteins. Nat Struct Mol Biol. 2005 Oct;12(10):842–6. doi: 10.1038/nsmb993.

29. Pękacz M, Basałaj K, Młocicki D, Kamaszewski M, Carretón E, Morchón R, Wiśniewski M, Zawistowska-Deniziak A. Molecular insights and antibody response to Dr20/22 in dogs naturally infected with *Dirofilaria repens*. Sci Rep. 2024 Jun 5;14(1):12979. doi: 10.1038/s41598-024-63523-9. Erratum in: Sci Rep. 2024 Jun 24;14(1):14499. doi: 10.1038/s41598-024-65401-w.

30. Wilbers RHP, Schneiter R, Holterman MHM, Drurey C, Smant G, Asojo OA, Maizels RM, Lozano-Torres JL. Secreted venom allergen-like proteins of helminths: Conserved modulators of host responses in animals and plants. PLoS Pathog. 2018 Oct 18;14(10):e1007300. doi: 10.1371/journal.ppat.1007300.

31. Darwiche R, Lugo F, Drurey C, Varossieau K, Smant G, Wilbers RHP, Maizels RM, Schneiter R, Asojo OA. Crystal structure of *Brugia malayi* venom allergen-like protein-1 (BmVAL-1), a vaccine candidate for lymphatic filariasis. Int J Parasitol. 2018 Apr;48(5):371–378. doi: 10.1016/j.ijpara.2017.12.003.

32. Perry, S. V. Vertebrate tropomyosin: distribution, properties and function. J. Muscle Res. Cell Motil. 22, 5–49 (2001).

33. Pérez-Morales D, Espinoza B. The role of small heat shock proteins in parasites. Cell Stress Chaperones. 2015 Sep;20(5):767–80. doi: 10.1007/s12192-015-0607-y.

34. Basha E, O’Neill H, Vierling E. Small heat shock proteins and α-crystallins: dynamic proteins with flexible functions. Trends Biochem Sci. 2012 Mar;37(3):106–17. doi: 10.1016/j.tibs.2011.11.005.

35. Dakshinamoorthy G, Samykutty AK, Munirathinam G, Shinde GB, Nutman T, Reddy MV, Kalyanasundaram R. Biochemical characterization and evaluation of a *Brugia malayi* small heat shock protein as a vaccine against lymphatic filariasis. PLoS One. 2012;7(4):e34077. doi: 10.1371/journal.pone.0034077. Epub 2012 Apr 5.

36. Gnanasekar M, Anandharaman V, Anand SB, Nutman TB, Ramaswamy K. A novel small heat shock protein 12.6 (HSP12.6) from *Brugia malayi* functions as a human IL-10 receptor binding protein. Mol Biochem Parasitol. 2008 Jun;159(2):98–103. doi: 10.1016/j.molbiopara.2008.02.010.

37. Massa C, Melani C, Colombo MP. Chaperon and adjuvant activity of hsp70: different natural killer requirement for cross-priming of chaperoned and bystander antigens. Cancer Res. 2005 Sep 1;65(17):7942–9. doi: 10.1158/0008-5472.CAN-05-0377.

38. Gandasegui J, Power RI, Curry E, Lau DC, O’Neill CM, Wolstenholme A, Prichard R, Šlapeta J, Doyle SR. Genome structure and population genomics of the canine heartworm *Dirofilaria immitis*. Int J Parasitol. 2024 Feb;54(2):89–98. doi: 10.1016/j.ijpara.2023.07.006.

39. Gregory WF, Atmadja AK, Allen JE, Maizels RM. The abundant larval transcript-1 and -2 genes of *Brugia malayi* encode stage-specific candidate vaccine antigens for filariasis. Infect Immun. 2000 Jul;68(7):4174–9. doi: 10.1128/IAI.68.7.4174-4179.2000.

40. Allen JE, Daub J, Guiliano D, McDonnell A, Lizotte-Waniewski M, Taylor DW, Blaxter M. Analysis of genes expressed at the infective larval stage validates the utility of *Litomosoides sigmodontis* as a murine model for filarial vaccine development. Infect Immun. 2000 Sep;68(9):5454–8. doi: 10.1128/IAI.68.9.5454-5458.2000.

41. Ali F, Brown A, Stanssens P, Timothy LM, Soule HR, Pritchard DI. Vaccination with neutrophil inhibitory factor reduces the fecundity of the hookworm *Ancylostoma ceylanicum*. Parasite Immunol. 2001 May;23(5):237–49. doi: 10.1046/j.1365-3024.2001.00383.x. PMID: 11309134.

42. Bower MA, Constant SL, Mendez S. *Necator americanus*: the Na-ASP-2 protein secreted by the infective larvae induces neutrophil recruitment in vivo and in vitro. Exp Parasitol. 2008 Apr;118(4):569–75. doi: 10.1016/j.exppara.2007.11.014.

43. Anand SB, Gnanasekar M, Thangadurai M, Prabhu PR, Kaliraj P, Ramaswamy K. Immune response studies with *Wuchereria bancrofti* vespid allergen homologue (WbVAH) in human lymphatic filariasis. Parasitol Res. 2007 Sep;101(4):981–8. doi: 10.1007/s00436-007-0571-2.

44. Wilbers RHP, Schneiter R, Holterman MHM, Drurey C, Smant G, Asojo OA, Maizels RM, Lozano-Torres JL. Secreted venom allergen-like proteins of helminths: Conserved modulators of host responses in animals and plants. PLoS Pathog. 2018 Oct 18;14(10):e1007300. doi: 10.1371/journal.ppat.1007300.

45. Chalmers IW, Hoffmann KF. Platyhelminth Venom Allergen-Like (VAL) proteins: revealing structural diversity, class-specific features and biological associations across the phylum. Parasitology. 2012 Sep;139(10):1231–45. doi: 10.1017/S0031182012000704.

46. Hartmann S, Sereda MJ, Sollwedel A, Kalinna B, Lucius R. A nematode allergen elicits protection against challenge infection under specific conditions. Vaccine. 2006 Apr 24;24(17):3581–90. doi: 10.1016/j.vaccine.2006.01.064.

47. Acosta-Benites J, Jara LM, Verastegui Pimentel M, Obregón Maldonado PM, Altamirano-Zevallos F, Valencia Mamani N, Gavidia CM. Development of the antigenic recombinant tropomyosin of *Echinococcus granulosus* in a bacterial system as a vaccinal candidate against canine echinococcosis. Rev Peru Med Exp Salud Publica. 2025 Jan 31;41(4):411–416. doi: 10.17843/rpmesp.2024.414.13854.

48. Jenkins RE, Taylor MJ, Gilvary NJ, Bianco AE. Tropomyosin implicated in host protective responses to microfilariae in onchocerciasis. Proc Natl Acad Sci U S A. 1998 Jun 23;95(13):7550–5. doi: 10.1073/pnas.95.13.7550.

49. Ehsan M, Haseeb M, Hu R, Ali H, Memon MA, Yan R, Xu L, Song X, Zhu X, Li X. Tropomyosin: An Excretory/Secretory Protein from *Haemonchus contortus* Mediates the Immuno-Suppressive Potential of Goat Peripheral Blood Mononuclear Cells In Vitro. Vaccines (Basel). 2020 Mar 1;8(1):109. doi: 10.3390/vaccines8010109. PMID: 32121527.

50. Yin Y, Martin J, McCarter JP, Clifton SW, Wilson RK, Mitreva M. Identification and analysis of genes expressed in the adult filarial parasitic nematode *Dirofilaria immitis*. Int J Parasitol. 2006 Jun;36(7):829–39. doi: 10.1016/j.ijpara.2006.03.002.

51. Saravanan N, Gray SA, Davis J, Puff-Carter CM, Kim J, Khatri V, Chauhan N, Carter D, Kalyanasundaram R. A next-generation human lymphatic filariasis vaccine candidate, rBmHAXT, for clinical development. NPJ Vaccines. 2026 Jul 8;11(1):171. doi: 10.1038/s41541-026-01497-7.

52. Chauhan N, Khatri V, Banerjee P, Kalyanasundaram R. Evaluating the Vaccine Potential of a Tetravalent Fusion Protein (rBmHAXT) Vaccine Antigen Against Lymphatic Filariasis in a Mouse Model. Front Immunol. 2018 Jul 2;9:1520. doi: 10.3389/fimmu.2018.01520.

53. Gray CA, Lawrence RA. A role for antibody and Fc receptor in the clearance of *Brugia malayi* microfilariae. Eur J Immunol. 2002 Apr;32(4):1114–20. doi: 10.1002/1521-4141(200204)32:4<1114::AID-IMMU1114>3.0.CO;2-B.

54. Mejia JS, Carlow CK. An analysis of the humoral immune response of dogs following vaccination with irradiated infective larvae of *Dirofilaria immitis*. Parasite Immunol. 1994 Mar;16(3):157–64. doi: 10.1111/j.1365-3024.1994.tb00335.x. PMID: 8208588.

55. 55. Morris Animal Foundation. https://www.morrisanimalfoundation.org/article/heartworm-dogs-cats.

56. Dagley JL, DiCosty U, Fricks C, Mansour A, McCall S, McCall JW, Taylor MJ, Turner JD. Current status of immunodeficient mouse models as substitutes to reduce cat and dog use in heartworm preclinical research. F1000Res. 2024 Sep 11;13:484. doi: 10.12688/f1000research.149854.2.56. Newport GR. Heat shock proteins as vaccine candidates. Semin Immunol. 1991 Jan;3(1):17-24. PMID: 1893121.

57. Khatri V, Chauhan N, Vishnoi K, von Gegerfelt A, Gittens C, Kalyanasundaram R. Prospects of developing a prophylactic vaccine against human lymphatic filariasis - evaluation of protection in non-human primates. Int J Parasitol. 2018 Aug;48(9-10):773–783. doi: 10.1016/j.ijpara.2018.04.002. Epub 2018 Jun 6. Erratum in: Int J Parasitol. 2018 Nov;48(13):1071. doi: 10.1016/j.ijpara.2018.09.001.

58. de Souza DK, Bockarie MJ. Current perspectives in the epidemiology and control of lymphatic filariasis. Clin Microbiol Rev. 2025 Jun 12;38(2):e0012623. doi: 10.1128/cmr.00126-23.

59. Infante-Duarte, C., Kamradt, T. Th1/Th2 balance in infection. Springer Semin Immunopathol 21, 317–338 (1999). 10.1007/BF00812260.

